# DNA METHYLATION PROFILING OF A CNIDARIAN-ALGAL SYMBIOSIS USING NANOPORE SEQUENCING

**DOI:** 10.1101/2021.02.01.429278

**Authors:** James L. Dimond, Nhung Nguyen, Steven B. Roberts

## Abstract

Symbiosis with protists is common among cnidarians such as corals and sea anemones, and is associated with homeostatic and phenotypic changes in the host that could have epigenetic underpinnings, such as methylation of CpG dinucleotides. We leveraged the sensitivity to base modifications of nanopore sequencing to probe the effect of symbiosis with the chlorophyte *Elliptochloris marina* on methylation in the sea anemone *Anthopleura elegantissima*. We first validated the approach by comparison of nanopore-derived methylation levels with CpG depletion analysis of a published transcriptome, finding that high methylation levels are associated with CpG depletion as expected. Next, using reads generated exclusively from aposymbiotic anemones, a largely complete draft genome comprising 243 Mb was assembled. Reads from aposymbiotic and symbiotic sea anemones were then mapped to this genome and assessed for methylation using the program Nanopolish, which detects signal disruptions from base modifications as they pass through the nanopore. Based on assessment of 452,841 CpGs for which there was adequate read coverage (approximately 8% of the CpGs in the genome), symbiosis with *E. marina* was, surprisingly, associated with only subtle changes in the host methylome. However, we did identify one extended genomic region with consistently higher methylation among symbiotic individuals. The region was associated with a DNA polymerase zeta that is noted for its role in translesion synthesis, which opens interesting questions about the biology of this symbiosis. Our study highlights the power and relative simplicity of nanopore sequencing for studies of nucleic acid base modifications in non-model species.

## INTRODUCTION

Endosymbiosis with algae is a widespread phenomenon among cnidarians inhabiting shallow tropical and temperate waters. The cnidarian-algal symbiosis is most notable as the trophic foundation of coral reef ecosystems, and plays a key role in the biology of scleractinian reef-building corals. For the host cnidarian, symbiosis can afford considerable nutritional benefits in the form of photosynthetically-fixed carbon that is translocated from the symbionts to the host cells in which they reside (Davy *et al.* 2012). Not surprisingly, symbiosis has physiological consequences for the host, many of which have been recently elucidated by studies of symbiosis-induced changes in host gene and protein expression (Rodriguez-Lanetty *et al.* 2006; Lehnert *et al.* 2014; Ganot *et al.* 2011; Oakley *et al.* 2016). These studies, involving a handful of facultatively symbiotic cnidarians capable of living naturally with and without symbionts, have revealed numerous affected pathways, including metabolism, membrane transport, cell adhesion, cell proliferation, cell-cell recognition, apoptosis, and oxidative stress (Rodriguez-Lanetty *et al.* 2006; Lehnert *et al.* 2014; Ganot *et al.* 2011; Oakley *et al.* 2016). Thus, symbiosis is associated with widespread homeostatic and phenotypic changes in the host (Rodriguez-Lanetty *et al.* 2006; Lehnert *et al.* 2014), yet the processes ultimately governing these changes at the molecular level are not well understood.

Epigenetic mechanisms are often implicated as regulators of homeostasis and phenotype, and include a suite of processes that act together to modify the genome without changing base sequences. To date, only two studies have investigated epigenetics in relation to the cnidarian-algal symbiosis. In *Aiptasia* sea anemones, Baumgarten *et al.* (2018) reported that microRNA (miRNAs) levels differed between symbiotic and aposymbiotic hosts, and differentially expressed miRNAs were associated with mRNA targets previously identified in studies of symbiosis-associated gene expression. Also in *Aiptasia,* Li *et al.* (2018) found differential DNA methylation levels according to symbiosis, and concluded that DNA methylation may play a role in maintaining transcriptional homeostasis under different symbiotic states.

DNA methylation, which most commonly involves conversion of cytosine to 5-methylcytosine, has traditionally been the most heavily researched and relatively well-understood epigenetic modification (Duncan *et al.* 2014). DNA methylation in invertebrate genomes is relatively sparse, yet it occurs most prominently in actively transcribed genes, where it is termed gene body methylation (Zemach *et al.* 2010; Sarda *et al.* 2012). While the function of gene body methylation remains a subject of study and debate, perhaps the most compelling evidence to date suggests that it plays a role in transcriptional fidelity by reducing spurious transcription (Neri *et al.* 2017). Because the majority of gene body methylation tends to occur in highly conserved housekeeping genes essential for cellular function (Sarda *et al.* 2012; Dimond & Roberts 2016), this supports the hypothesis that methylation is particularly important where transcriptional fidelity is essential. Indeed, the study by Li *et al.* (2018) on *Aiptasia* supported the hypothesis that DNA methylation reduces spurious transcription.

Continued advances in genomic sequencing technologies are allowing ever-greater resolution of methylomes. With the advent of nanopore sequencing, in which the base composition of full-length nucleic acid molecules passing through a nanopore is quantified using electrical currents, base modifications such as DNA methylation can be readily detected without additional library preparation steps (Jain *et al.* 2016; Simpson *et al.* 2017). Moreover, the ability to sequence full-length molecules permits construction of reference genomes with lower throughput and cost than short read technologies. Together, these properties make nanopore sequencing a promising tool for epigenetics research on non-model species.

In this study, our goal was to test the efficacy of methylation profiling using nanopore sequencing, and use it to determine if symbiosis is associated with differential methylation in *Anthopleura elegantissima* sea anemones. A common member of intertidal communities along the Pacific coast of North America, *A. elegantissima* is unique in its ability to exist either aposymbiotic or symbiotic with one (and occasionally both) of two distinct algal symbionts: the chlorophyte *Elliptochloris marina* or the dinoflagellate *Brevolium muscatinei* (Dimond *et al.* 2011). This facultative symbiosis makes *A. elegantissima* an excellent model system for research on cnidarian-algal symbiosis.

## METHODS

### Specimen collection

At Lawrence Point, Orcas Island, Washington, USA, eight *A. elegantissima* were collected within an approximately 15 cm x 15 cm area from the underside of an intertidal boulder in June 2018. The individuals were chosen from within this small area because they likely represented a single aggregated clone, and yet individuals in two different symbiotic states could be seen. *Anthopleura elegantissima* is well known for its aggression towards genetically unrelated clones (Ayre & Grosberg 1995), and clonal individuals can therefore be selected by targeting individuals from within a single aggregation. Four aposymbiotic individuals and four symbiotic individuals hosting the chlorophyte *E. marina* were chosen; the symbiotic status of these sea anemones can be easily discerned in the field based on color alone, since aposymbiotic individuals appear white and symbiotic animals are green (Figure 1A). This sampling scheme was chosen to select two clearly different, naturally occurring phenotypes while minimizing environmental and genetic effects. Specimens were flash frozen in liquid nitrogen within one hour of collection. Just prior to freezing, tentacles excised from anemones were used to confirm the absence of *E. marina* in aposymbiotic animals using epifluorescence microscopy; red fluorescence from the chlorophyll-containing *E. marina* makes their presence readily apparent (Figure 1B).

**Figure 1.**
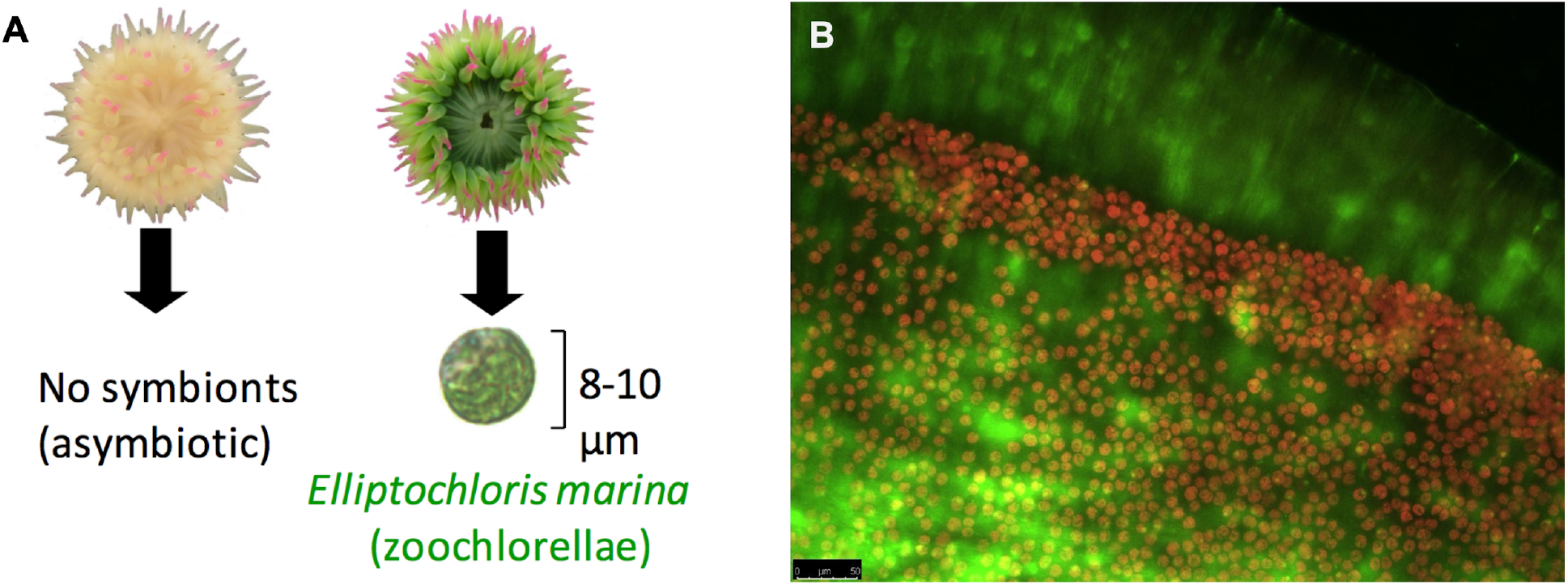
(A) Symbiotic and aposymbiotic *A. elegantissima* are easily discernable based on the green color imparted by the *E. marina* symbionts. (B) The presence/absence of symbionts is verified in an excised tentacle viewed under epifuorescence microscopy, where symbiont red fluorescence makes them highly visible. Green fluorescence is due to green fluorescent protein (GFP) of the host. Scale bar = 50 μm.

### DNA sequencing

Frozen sea anemone tissue was ground to a fine powder with a mortar and pestle in liquid nitrogen prior to genomic DNA extraction with the Qiagen DNeasy Blood and Tissue kit. The Qubit BR assay was used to assess DNA quantity, followed by 1% agarose gel electrophoresis to confirm the presence of high molecular weight DNA (≥10 kb). Four DNA libraries were prepared for sequencing with the Oxford Nanopore Technologies (ONT) MinION platform. The MinION was connected to an Apple iMac with 1TB SSD, 32 GB RAM, 3.5 GHz Intel Core i7, and running MacOS 10.12.5 and MinKNOW 1.15.4. The first library was prepared with gDNA from a single aposymbiotic individual using the PCR-free, transposase-based ONT Rapid Sequencing Kit (SQK-RAD004) following manufacturer guidelines. Sequencing was done on two FLO-MIN 106D R9.4 flow cells.

A second round of sequencing runs was performed several months later. Three libraries with one aposymbiotic individual and one symbiotic individual each, both individually barcoded, were prepared using the PCR-free ONT Ligation Sequencing Kit with the Native Barcoding Expansion Kit (SQK-LSK109 and EXP-NBD103), following the manufacturer’s 1D native barcoding gDNA protocol. Each library was run on a separate FLO-MIN 106D R9.4 flow cell. Basecalling and demultiplexing was performed with ONT Albacore Sequencing Pipeline Software v. 2.3.3, and sequencing adapters were removed with Porechop v. 0.2.4 (Wick *et al.* 2017).

### Comparison of nanopore-derived methylation with CpG depletion estimates

Raw reads from the first round of sequencing were mapped to a published *A. elegantissima* transcriptome (Kitchen *et al.* 2015) using minimap2 v. 2.14 (Li 2018). DNA methylation patterns for mapped reads were analyzed with Nanopolish v. 0.10.1 (Simpson *et al.* 2017). Nanopolish analyzes CpG motifs in short (11-34 bp) k-mer sequences and distinguishes 5-methylcytosine from unmethylated cytosine based on signal disruptions in raw ONT FAST5 sequence data. The program calculates log-likelihood ratios for base modifications of each read, with positive values indicating support for modification (Figure S1). Nanopolish output was summarized using the helper script calculate_methylation_frequency.py (Simpson *et al.* 2017), which summarizes methylation frequency by genomic coordinates. Nanopolish methylation calls were then subject to filtering using a minimum read count of 10 reads per CpG motif. Further filtering excluded transcriptome sequences with methylation called for fewer than 20% of CpGs in the sequence.

To estimate CpG depletion--an indicator of evolutionary-scale methylation patterns--in the *A. elegantissima* transcriptome, the number of CpGs observed vs. expected, CpG O/E, was calculated as:

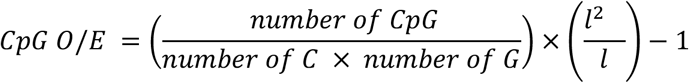

where *l* is the length of the sequence. Outliers were excluded by setting a CpG O/E threshold of > 0.001 and < 2.

### *A. elegantissima* draft genome generation

Using reads from only aposymbiotic individuals from both rounds of sequencing, a draft *A. elegantissima* genome was generated with wtdbg2, a de novo sequence assembly program designed for long noisy reads such as those produced by ONT sequencing (Ruan & Li 2020). Wtdbg2 first assembles raw reads without error correction, then builds a consensus from intermediate assembly output. ONT settings with default values were used.

### Methylation analysis of symbiotic and aposymbiotic sea anemones

DNA methylation patterns for each sequenced *A. elegantissima* individual were analyzed with Nanopolish v. 0.10.1 (Simpson *et al.* 2017) as described above. The draft *A. elegantissima* genome was used by Nanopolish for sequence mapping with minimap2 v. 2.14 (Li 2018). A threshold of at least 3 called sites per k-mer per sample was set to exclude low-coverage data. To investigate methylation patterns at individual genomic regions, pairwise t-tests were computed at each genomic position. The Benjamini-Hochberg procedure was applied to t-test p-values for false discovery rate (FDR) correction at ⍺ = 0.05. The Python tool methplotlib v. 0.1.1 (De Coster *et al.* 2020) was used to visualize regions of interest graphically and to further summarize data.

### Data availability

A repository containing datasets, notebooks, scripts, and output files is publicly available at https://github.com/jldimond/Ae_meth. Upon manuscript acceptance, these files will be published in an online repository. Additionally, sequencing reads will be deposited in the NCBI Sequence Read Archive.

## RESULTS

### Sequencing

Sequencing statistics are shown in Table 1. The ONT Rapid DNA sequencing protocol, in which a single aposymbiotic individual was sequenced, produced a total of 3.8 Gb from two flow cell runs. Mean read length from Rapid sequencing libraries (1,260 bp) was relatively short by comparison with read length obtained from the ONT Ligation DNA sequencing protocol (overall mean 4,177 bp). Ligation sequencing libraries produced an average of 688 Mb per sample, with a maximum read length of 72,055 bp.

**Table 1.** Sequencing statistics by sample. Most samples were sequenced in groups of two, and this is reflected in the library number. All libraries except the Rapid library were run on one flow cell each. The statistics reflect sequences that passed quality filtering only, and only barcoded sequences are reported for Ligation libraries. The prefix ‘Apo’ under the sample category denotes an aposymbiotic sea anemone, whereas ‘Sym’ denotes a symbiotic anemone.

| **Sample** | **Library** | **Total reads** | **Total base pairs** | **Mean bp** | **Max bp** | **N50 bp** |
| --- | --- | --- | --- | --- | --- | --- |
| Apo Rapid | Rapid (2 flow cells) | 3,044,962 | 3,835,969,017 | 1,260 | 48,696 | 1,753 |
| Apo 1 | Ligation #1 | 105,499 | 494,824,366 | 4,711 | 54,995 | 7,668 |
| Sym 1 | Ligation #1 | 139,157 | 669,243,165 | 4,765 | 70,128 | 8,465 |
| Apo 2 | Ligation #2 | 150,068 | 647,455,529 | 4,314 | 50,946 | 7,252 |
| Sym 2 | Ligation #2 | 347,615 | 1,208,181,769 | 3,476 | 72,055 | 6,373 |
| Apo 3 | Ligation #3 | 153,161 | 632,263,257 | 4,128 | 54,707 | 6,581 |
| Sym 3 | Ligation #3 | 130,673 | 479,395,377 | 3,669 | 65,555 | 7,048 |

### CpG depletion and methylation analysis

The transcriptome from Kitchen et al. (2015) represented a total of 142,933 sequences. After a coverage threshold was set for 10 nanopore reads per CpG motif, 66,734 transcriptome contigs had associated nanopore reads mapped. After the data were further processed with a threshold of 20% coverage of transcriptome data by nanopore data, 50,601 transcriptome contigs had associated nanopore methylation data. Thus, slightly less than one-third of the transcriptome was analyzed for CpG depletion and nanopore-derived methylation patterns.

The assessment of CpG depletion revealed that the vast majority of transcriptome sequences have high CpG O/E, reflecting low levels of evolutionary-scale DNA methylation frequency as is common among invertebrate genomes (Figure 2A). There was an inverse relationship of CpG O/E to the nanopore-derived methylation frequency. Comparison of fully unmethylated to fully methylated transcripts further illustrates much higher CpG O/E among fully unmethylated transcripts (t-test, p < 0.001; Figure 2B).

**Figure 2.**
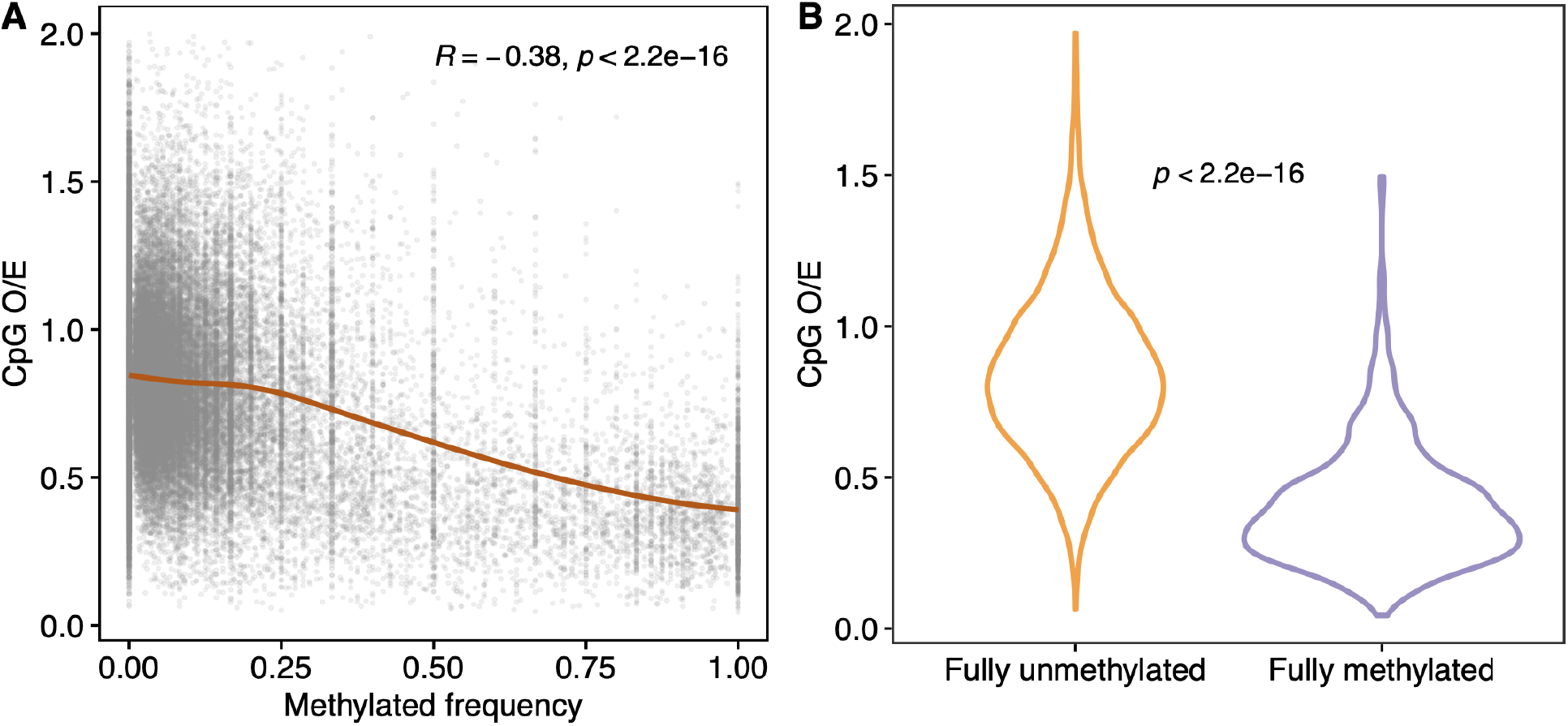
(A) Relationship between nanopore-derived methylation frequency of reads mapped to the *A. elegantissima* transcriptome and CpG depletion estimates (CpG O/E) of those sequences. A LOESS (locally estimated scatterplot smoothing) moving average is fitted to the data, with pearson correlation results shown in upper margin. (B) Violin plot showing CpG depletion estimates for sequences determined to be either fully unmethylated (methylation frequency = 0) or fully methylated (methylation frequency = 1) via nanopore sequencing. T-test results are shown.

### Draft genome

The wtdbg2-generated draft genome comprised 243 Mb, including 5359 contigs with an N50 of 87 kb and N90 of 19.2 kb. All aposymbiotic sequences were used to generate the draft genome, providing a total of 5.6 Gb and an estimated coverage of 23x.

### Methylation analysis

After filtering for minimum read coverage, a total of 452,841 CpGs in 171,545 k-mers were analyzed for methylation, accounting for 7.8% of the ~5,818,825 CpGs in the *A. elegantissima* draft genome. Because Nanopolish cannot resolve methylation of individual CpGs within ~6 bases each other, k-mers often included more than one CpG (mean = 2.6 CpGs per k-mer). Overall CpG methylation frequency was very low; mean methylation frequency was 0.055 (S.D. = 0.002) for aposymbiotic anemones and 0.059 (S.D. = 0.001) for symbiotic anemones. While there was a clear trend of higher methylation among symbiotic individuals, the difference was not significant (t-test, p = 0.057; Figure 3A). Similarly, principal components analysis of methylation patterns also did not reveal any strong differences between symbiotic states, yet there was some separation of aposymbiotic and symbiotic individuals along PC2, representing 9.4% of the variance (Figure 3B).

**Figure 3.**
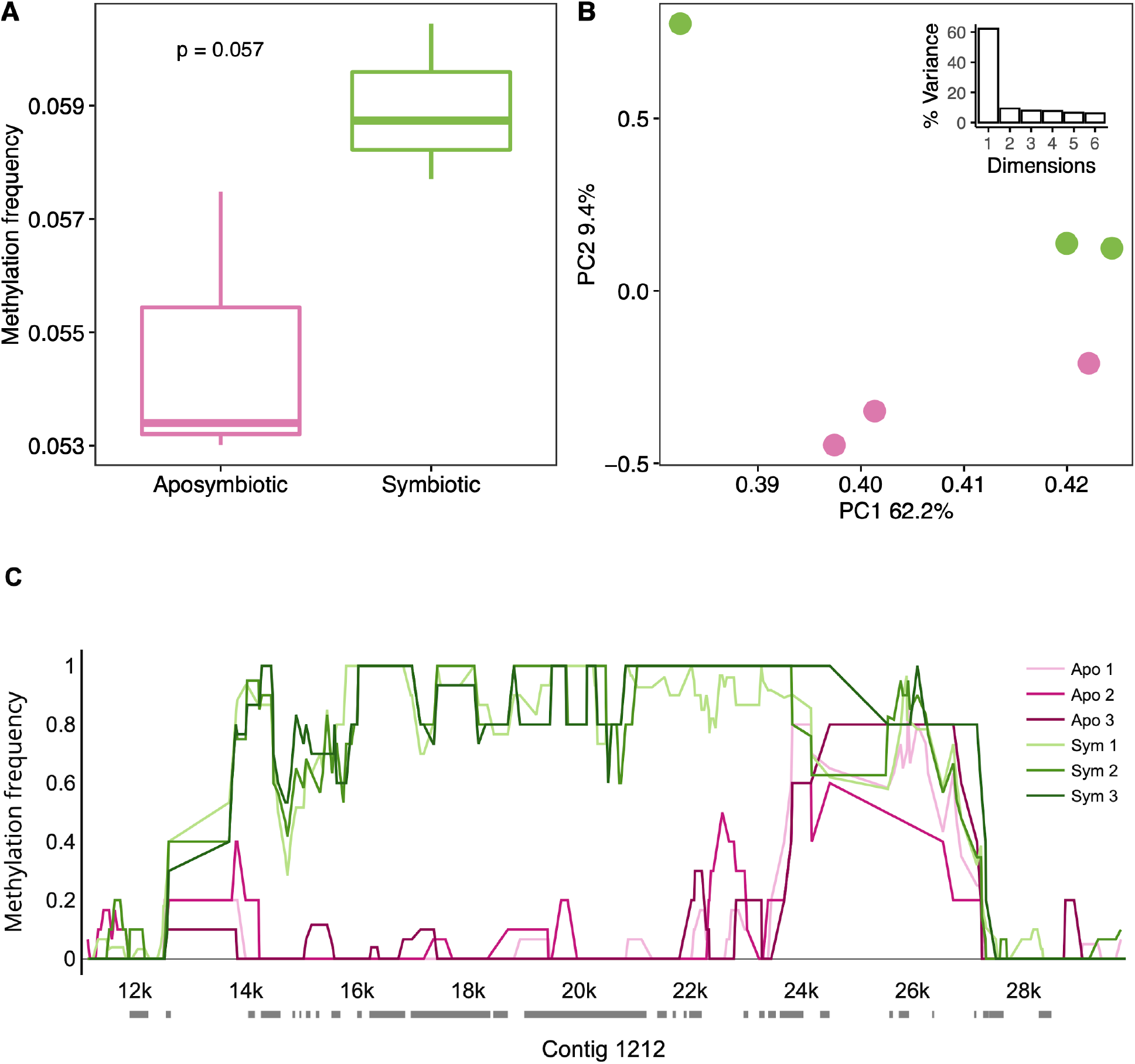
Comparative methylation frequency among aposymbiotic and symbiotic *A. elegantissima*. (A) Boxplots of genomewide methylation frequency of aposymbiotic and symbiotic specimens. Horizontal lines represent the median, with box edges representing interquartile range, and whiskers 1.5 times the interquartile range. T-test results are shown in the upper left. (B) Principal components analysis of methylation frequency, showing individuals plotted by the first two principal components. Component loadings are shown in the axis labels and in the inset scree plot. Symbiotic state is represented by colors identical to (A). (C) Methylation frequency patterns associated with a 20 kb region of contig 1212 in aposymbiotic and symbiotic *A. elegantissima*. Aposymbiotic specimens are represented by pink hues while symbiotic individuals are shown in green. Transcript alignments from Kitchen et al. (2015) are shown as horizontal gray bars along the x-axis.

At each genomic position, none of the t-tests passed FDR correction. However, there were 9 comparisons that were “perfect” differences with no variance; t-tests are not possible with these data and the p-value is hypothetically infinite. We examined these regions and found a particularly compelling differentially methylated region on contig 1212, in which an approximately 10 kb region showed consistently high methylation frequency among symbiotic individuals, while methylation frequency among aposymbiotic individuals was near zero (Figure 3C; Figure S2). Twenty-one transcripts from (Kitchen *et al.* 2015) were mapped to this region. While eight of these transcripts were either uncharacterized or without hits, the remaining 13 were interspersed along the entire length of the differentially methylated region and were all associated with a DNA polymerase zeta catalytic subunit. This was confirmed by NCBI megablast query of the 20 kb region of contig 1212 shown in Fig. 3C, which returned best matches with other sea anemone mRNA sequences designated as DNA polymerase zetas (Figure S3).

## DISCUSSION

Nanopore sequencing on the ONT MinION platform combined with the Nanopolish methylation detection program (Simpson *et al.* 2017) provided high-quality DNA methylation data at near-base resolution. We were also able to easily generate an *A. elegantissima* draft genome with which to map reads. At 243 Mb, our *A. elegantissima* assembly is comparable in size to those of other actiniarians, including *Nematostella vectensis* (329 Mb; Putnam *et al.* 2007) and *Aiptasia pallida* (260 Mb; Baumgarten *et al.* 2015), as well as closely-related scleractinian corals (234-653 Mb; Shinzato *et al.* 2021). Based on the 23x coverage estimate as well as very similar genome size estimates from a concurrent short-read *A. elegantissima* genome effort (Holland Elder, pers. comm.), the draft genome generated here is likely to be largely complete. While a fully annotated, hybrid genome using both data sources is forthcoming, the basic assembly generated here provided a highly efficient and effective means of isolating host sequences from symbiont sequences, which is an initial hurdle for functional genomics work on symbiotic organisms.

As expected, a large portion of the *A. elegantissima* transcriptome has relatively high CpG O/E, reflecting the low levels of DNA methylation typical among invertebrates at evolutionary timescales (Sarda *et al.* 2012). Numerous studies have identified the small fraction of genes that tend to be highly methylated in invertebrates as conserved housekeeping genes (Gavery & Roberts 2010; Sarda *et al.* 2012; Dixon *et al.* 2014; Dimond & Roberts 2016). When comparing CpG O/E content to methylation frequency derived from nanopore sequencing, a negative correlation was found. This inverse relationship mimics those found in other invertebrates when comparing CpG O/E to actual measures of methylation (Sarda et al. 2012; Dixon *et al.* 2016). Thus, CpG O/E corroborates the nanopore-derived methylation frequency data reported here.

Surprisingly, methylation levels and profiles did not differ greatly between aposymbiotic and symbiotic anemones hosting *E. marina.* This was unexpected, since Li *et al.* (2018) found more pronounced differences between symbiotic and aposymbiotic *E. pallida*. In their study, albeit with much greater genomic coverage (53x per individual) than our study, Li *et al.* (2018) found over two thousand genes (approximately 7.3% of *E. pallida* genes) exhibiting significant methylation differences according to symbiotic state. Aside from lower genomic coverage in our study, two other factors may have contributed to the relatively subtle effects of symbiosis observed here. First, differences in experimental design may have been a factor. Whereas our study was a mensurative study of anemones in their natural state, Li *et al.* (2018) performed experimental bleaching and reinfection of anemones with symbionts. Secondly, in addition to obvious differences in host taxa between studies, the dinoflagellate symbiont of *E. pallida* in Li *et al.* (2018) is very different from the chlorophyte symbiont of *A. elegantissima* in our study. It is possible that symbiosis with *E. marina* does not elicit significant changes in host methylation because of its relatively limited contributions to host nutrition (Bergschneider & Muller-Parker 2008), or because it is more effectively able to evade detection by the host’s immune system than dinoflagellate symbionts. For example, there is evidence that successful establishment of cnidarian symbiosis by highly compatible symbionts is associated with limited changes in host gene expression relative to unsuccessful establishment by less compatible symbionts (Voolstra *et al.* 2009).

Although there was no substantial remodeling of the host methylome associated with symbiosis, there was some evidence of differential methylation between aposymbiotic and symbiotic genomes. A particularly compelling differentially methylated region was located on contig 1212, where much higher methylation among symbiotic individuals was consistent for a stretch of approximately 10 kb. This region was noted for multiple transcripts associated with a DNA polymerase zeta catalytic subunit. DNA polymerase zeta is a translesion synthesis polymerase that is able to bypass DNA lesions and complete transcription of damaged DNA, albeit at the cost of potential mutations (Gan *et al.* 2008). In yeast, for example, DNA polymerase zeta is critical for error-free replication past thymine glycol, a common DNA lesion created by free radicals (Johnson *et al.* 2003). Given that oxidative stress is a prominent physiological consequence of hosting algal endosymbionts (Weis 2008), this raises intriguing possibilities about how symbiosis might influence this process. Since methylation is generally associated with active transcription (Li *et al.* 2018), one hypothesis is that higher methylation of this DNA polymerase zeta gene among symbiotic anemones would be associated with higher transcription. That being said, it cannot be assumed that higher methylation is associated with higher transcription or exon inclusion, as the relationship between methylation and transcription is nuanced (Jones 2012) and the inclusion of exons can be either enhanced or suppressed by DNA methylation in a context-specific manner (Yearim *et al.* 2015). Future work targeting both methylation and transcription in this region may lead to further insights.

Two recent studies with symbiotic cnidarians have found limited correlation between differentially expressed and differentially methylated genes. In their study of symbiosis in E. pallida, Li *et al.* (2018) found numerous differentially methylated and differentially expressed genes, yet very little overlap between the two sets of genes. Similarly, Dixon *et al.* (2018) found only weak correlation between changes in methylation and transcription at the individual gene level. Interestingly, however, both of these studies found associations between changes in methylation and transcription when considering gene functional groups or pathways rather than individual genes. Both studies concluded that there is some complementarity between methylation and transcription at a broad functional level, yet precise mechanisms remain a subject for further study.

### Conclusion

Nanopore sequencing is clearly a powerful genomics technology for investigations of DNA base modifications, and it is a strong alternative to other methods due to the relative ease of library preparation, sequencing, and draft genome generation. Many of these qualities make it a good choice for investigations of non-model species. Surprisingly, only subtle differences in methylation associated with *A. elegantissima*’s unique symbiosis with *E.marina* were observed. While the presence of *E. marina* does not appear to strongly influence the host methylome, a region with clear and consistent differential methylation between symbiotic states was identified. The potential involvement of DNA polymerase zeta and the role of translesion synthesis in this symbiosis deserves further attention, and may yield new insights into the cnidarian-algal symbiosis as well as the potential functions of DNA methylation.

---

**Figure S1.**
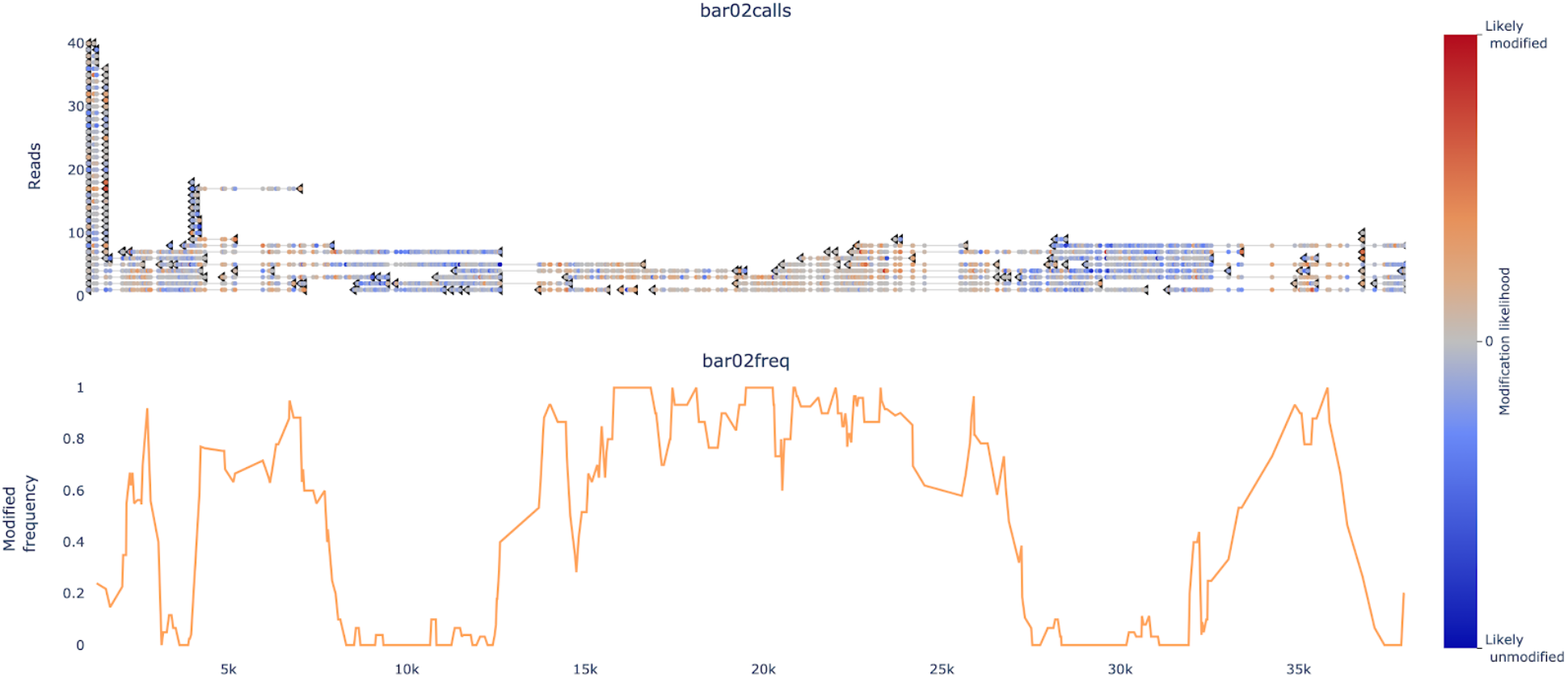
Example of methylation frequency estimation with Nanopolish (Simpson *et al.* 2017) along contig 1212. Upper panel shows individual reads colored by modification likelihood estimated by Nanopolish, with color ramp legend in right margin. Values above 0 indicate positive support for base modification/methylation. Lower panel shows modification frequency averaged from reads.

**Figure S2.**
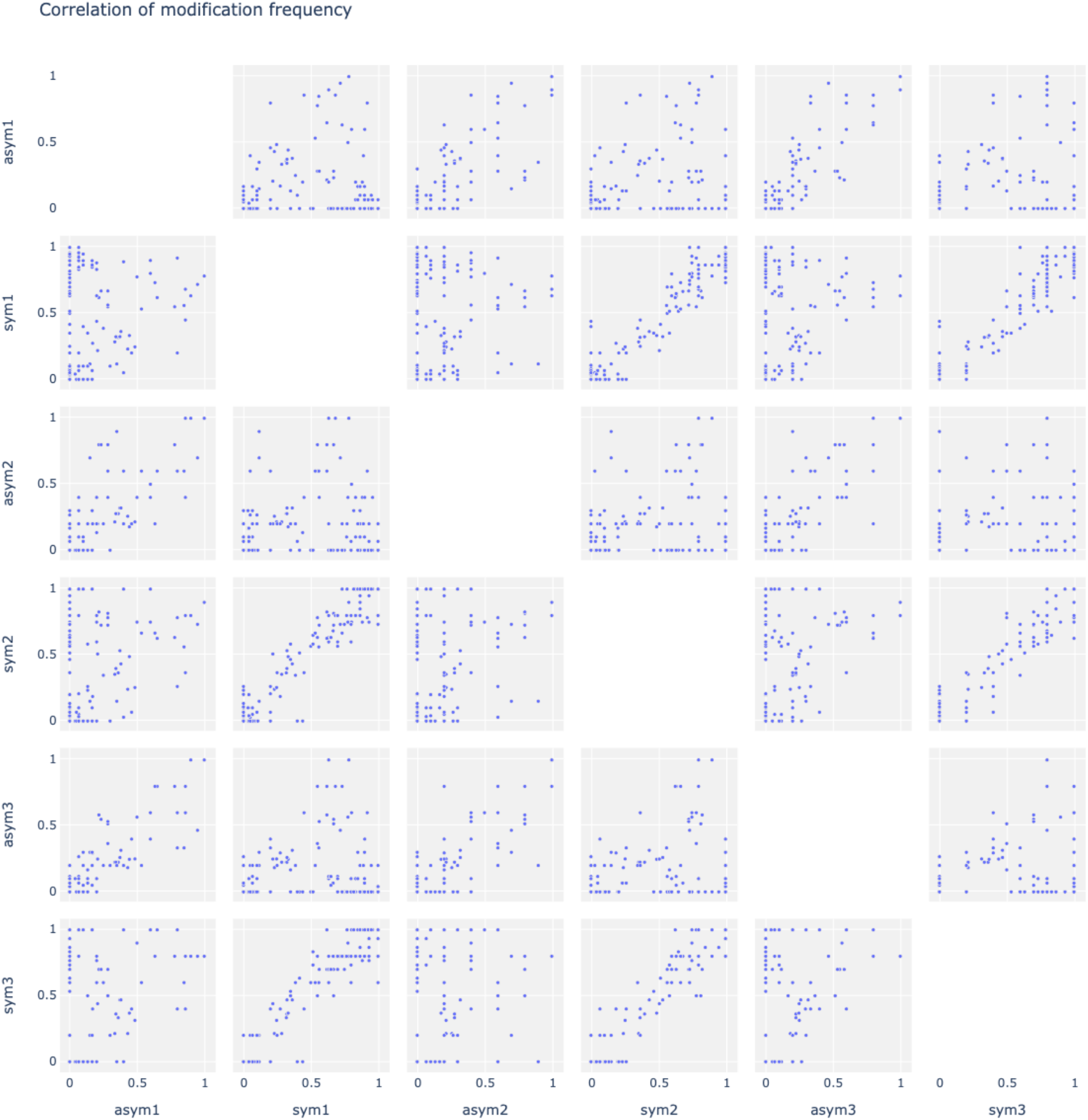
Correlation plots showing pairwise comparisons of modification frequency for each sample along the region of contig 1212 shown in Fig. 3C. Note that modification frequencies of symbiotic specimens (‘sym’) show strong positive correlations.

**Figure S3.**
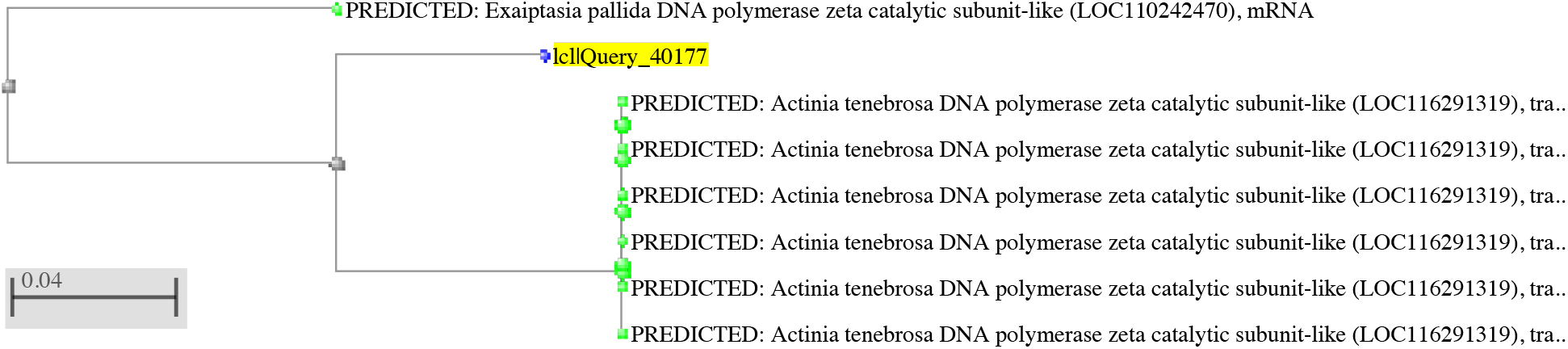
Neighbor-joining tree of NCBI megablast results with the 20 kb region of contig 1212 shown in Fig. 3C as the query sequence (highlighted in yellow). The closest matches were DNA polymerase zeta mRNA sequences from other sea anemone species.

